# The Charlson Index is insufficient to control for comorbidities in a national trauma registry

**DOI:** 10.1101/329581

**Authors:** Audrey Renson, Marc A. Bjurlin

## Abstract

**Background:** The Charlson Comorbidity Index (CCI) is frequently used to control for confounding by comorbidities in observational studies, but its performance as such has not been studied. We evaluated the performance of CCI and an alternative summary method, logistic principal component analysis (LPCA), to adjust for comorbidities, using as an example the association between insurance and mortality.

**Materials and Methods:** Using all admissions in the National Trauma Data Bank 2010-2015, we extracted mortality, payment method, and 36 ICD-9-derived comorbidities. We estimated ORs for the association between uninsured status and mortality before and after adjusting for CCI, LPCA, and separate covariates. We also calculated standardized mean differences (SMDs) of comorbidity variables before and after weighting the sample using inverse probability of treatment weights (IPTW) for CCI, LPCA, and separate covariates.

**Results:** In 4,936,880 admissions, most (68.3%) had at least one comorbidity. Considerable imbalance was observed in the unweighted sample (mean SMD=0.086, OR=1.17), which was almost entirely eliminated by IPTW on separate covariates (mean SMD=0.012, OR=1.36). The CCI performed similarly to the unweighted sample (mean SMD=0.080, OR=1.25), while 2 LPCA axes were better able to control for confounding (mean SMD=0.04, OR=1.31). Using covariate adjustment, the CCI accounted for 56.1% of observed confounding, whereas 2 LPCA axes accounted for 91.3%.

**Conclusions:** The use of the CCI to adjust for confounding may result in residual confounding, and alternative strategies should be considered. LPCA may be a viable alternative to adjusting for each comorbidity when samples are small or positivity assumptions are violated.

## Introduction

Large trauma registries have been a major source for developments in injury research, as they contain very large numbers of patients with relatively accurate information on patient health status, injury characteristics, and operative management.^1^ Despite these strengths, trauma registries are observational in nature, and studies of the effects of exposures on outcomes will often suffer from confounding, or nonrandom exposure assignment affected by variables which also impact the outcome of interest.

One category of confounding variables affecting many studies is comorbidities, or pre-existing diagnoses separate from the principal reason for admission, which naturally can impact mortality and other outcomes.^2^ Comorbidities in healthcare databases often present as many binary variables indicating presence or absence of a large number of conditions. This high dimensionality (i.e. presence of a large number of variables) of the data presents a problem for adjustment of confounding, as inclusion of all comorbidity variables in a regression model may not be feasible, due to multicollinearity problems, inadequate degrees of freedom, or violation of positivity (i.e. some categories of confounder-by-exposure crosstabs may contain zero observations).

A common approach to control for confounding is to use a comorbidity summary score, most commonly the Charlson Comorbidity Index (CCI). The CCI was developed in a single institution to predict 1-year mortality risk among patients admitted to a medical ward,^3^ and is considered a highly valid and reliable means to do so.^4^ The score is also widely used to control for confounding, whether by entering it as a covariate in a regression model or other similar means – a use which has not been properly validated. Studies purporting to investigate the utility of the CCI for confounding control have relied on measures of ability to predict the outcome (e.g., mortality).^5, 6^ The predictive ability of CCI and other comorbidity scores (e.g., using *C*-statistics) is frequently cited as relevant to assessing their role in “risk adjustment” and “casemix adjustment”,^7-11^ requiring the assumption that prediction of mortality equals control of confounding. However, this is not necessarily the case, as the degree of confounding conferred by a variable reflects its effect on both the exposure and outcome.^12^

One alternative is to perform dimension reduction, such as principal component analysis (PCA), which reduces large numbers of variables to a few uncorrelated variables capturing an arbitrarily large proportion of variance in the data. With a few considerations, these reduced variables can be entered as covariates into a regression model, a potentially efficient way to reduce confounding that eliminates concerns of multicollinearity, positivity, or degrees of freedom. PCA has previously been used to describe comorbidity patterns,^13^ however, its capacity to adequately control for confounding by comorbidity has not, to our knowledge, been tested.

The aim of this study was to evaluate and compare the ability of the CCI and a PCA-based method to adjust for measured confounding by pre-existing comorbidity variables in a large, national trauma registry. Results of this study can inform best practices for data analysis of this and other similar large healthcare databases.

## Materials and Methods

This was a retrospective cohort study including inpatient hospital data from the American College of Surgeons (ACS) National Trauma Data Bank (NTDB), the largest national repository on trauma admissions. The NTDB contains data on trauma-related hospital admissions from over 700 facilities, including both verified trauma centers and non-designated facilities. NTDB includes all patient admissions with International Classification of Disease, Ninth Revision, Clinical Modification (ICD-9-CM) discharge diagnosis 800.00-959.9, excluding late effects of injuries (905-909), superficial injuries (905-909), and foreign body cases (930-939). Patient information is provided to the NTDB on a voluntary basis by participating trauma centers, de-identified, maintained in a secure database, cleaned and standardized to ensure data quality.^14^ As an approved public dataset, the NTDB contains no patient identifiers and is not considered human subjects research. As a result IRB approval is not needed and was not obtained. The study sample was composed of all patient admissions reported to the NTDB between 2010 and 2015, excluding patients whose hospital charges were paid by worker’s compensation, no fault auto, not billed, or “other”.

### Measures

The NTDB does not directly measure insurance status, but does record mode of payment for hospital charges. Patients were classified as uninsured if the mode of payment was self-pay or unreported; otherwise they were classified as insured. We chose this exposure-outcome relationship as it has been frequently observed, and is very likely to be confounded by comorbid conditions – that is, patients with comorbid conditions have more of need for insurance, and also have more underlying risk of mortality, compared to otherwise healthy patients.

Comorbidities were recorded in the NTDB based on International Classification of Diseases, Ninth Revision, Clinical Modification (ICD-9-CM) diagnosis codes for following conditions: alcohol use disorder, bleeding disorder, currently receiving chemotherapy for cancer, congenital anomalies, congestive heart failure, current smoker, chronic renal failure, cerebrovascular accident, diabetes mellitus, disseminated cancer, advanced directive limiting care, history of angina within 30 days, history of myocardial infarction, history of peripheral vascular disease (PVD), hypertension requiring medication, prematurity, chronic obstructive pulmonary disorder (COPD), steroid use, cirrhosis, dementia, major psychiatric illness, drug use disorder, or attention deficit disorder/attention deficit hyperactivity disorder (ADD/ADHD), or other. For definitions of each of these conditions, see the ACS National Trauma Data Standard data dictionary.^15^

### Statistical Analysis

To summarize comorbid conditions, we calculated the CCI adapted to ICD-9 codes as previously described (shown in Table 1),^16^ as well as logistic principal components analysis (LPCA), an extension of conventional PCA to allow for binomially distributed variables, analogous to the distinction between linear and logistic regression.^17^ We selected the maximum number of LPCA axes to include in regression models based on visual identification of an inflexion point in the scree plot, or bar plot displaying the marginal variance explained by each additional axis.

**Table 1.** Charlson Comorbidity Index (CCI), as adapted for use with ICD-9-CM data by Deyo et al (1992)^16^.

| Points | Condition |
| --- | --- |
| 1 | Cerebrovascular disease |
| 1 | Chronic pulmonary disease |
| 1 | Congestive heart failure |
| 1 | Myocardial infarction |
| 1 | Peripheral vascular disease |
| 2 | Diabetes |
| 2 | Hemiplegia |
| 2 | Renal disease |
| 2 | Tumor without metastases |
| 3 | Liver disease |
| 6 | Metastatic solid tumor |

Under the counterfactual (a.k.a. potential outcomes) framework, in order for a causal effect to be reliably estimated from data, exposure groups must have *exchangeability* with respect to confounding variables.^18, 19^ In other words, the aim when controlling for confounders is to effect maximum covariate balance between exposure groups. A well-conducted randomized controlled trial is the gold standard for confounding control, in that covariate balance is achieved between the (randomly allocated) exposure groups, a condition which can be easily assessed by comparing distributions of observed confounders between the treatment arms.

In contrast, there is no clear metric of confounding control when using multivariable regression-based covariate adjustment, the most common means of adjustment in a non-randomized study.

However, pre-analytic methods of covariate adjustment, including propensity score stratification, matching, and inverse probability of treatment weighting (IPTW), have gained considerable favor over regression adjustment, in part because of their allowance for direct assessment of covariate balance. We used IPTW because it tends to perform well in simulations.^20-22^

To calculate IPTW weights, we used logistic regression models, regressing uninsured status on comorbidity variables. To determine the functional relationship of each confounder summary to the exposure, we examined plots of proportion insured against quantiles of the confounder summary, along with penalized smoothing splines. In the presence on nonlinearity, we included polynomial terms in a forward stepwise manner in the model to estimate the weights, as long as their inclusion reduced the Akaike’s Information Criterion of the model. This led to the inclusion of quadratic terms for all three LPCA axes and for CCI.

Weights were then calculated as the proportion uninsured in the sample, divided by the predicted probability of receiving the treatment that was actually received. As is common practice, weights were truncated at the 0.2^nd^ and 99.8^th^ percentiles, such that values exceeding the threshold value were set to that value.^23^ As a sensitivity analysis, we also conducted all described analyses with non-truncated weights, and results were identical.

To measure covariate balance under IPTW, we used the standardized mean difference (SMD), a.k.a. Cohen’s *d*,^24^ which in the case of continuous covariates represents the difference in mean divided by the pooled standard deviation, and is easily generalized to binary and categorical variables.^25^ Because all covariates were categorical, the SMD weighted by IPTW was calculated by a weighted chi-squared statistic transformed into SMD with the following formula^26^:

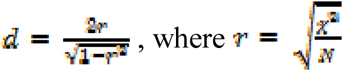

In this formula, *d* refers to the SMD, χ^2^ refers to the chi-squared statistic for the confounder against the exposure, and *N* refers to the sample size used to calculate this statistic. In addition to comparison of SMDs, to evaluate the performance of separate covariates, CCI, and LPCA, we also fit logistic regression models using both IPTW and covariate adjustment, with separate models adjusted for the CCI, up to the first three principal component scores, and each comorbid condition separately. We considered the models adjusted for separate covariates to be the gold standard, against which we compared both CCI and LPCA, in terms of both the estimated SMD and the value of the odds ratio for mortality in uninsured vs. insured patients.

Finally, we calculated the total proportion of observed confounding accounted for by each method, defined as the difference in log odds ratio between the adjusted (CCI or LPCA) and crude, divided by the difference in log odds ratio between the adjusted (separate covariates) and crude.

All statistical analyses were conducted in R software version 3.4.2 for Windows,^27^ using the survey package^28, 29^ to calculate weighted Chi squared statistics, the logisticPCA^30^ package for logistic principal component analysis, and the compute.es package^31^ to calculate SMDs.

## Results

Between 2010 and 2015, there were 4,936,880 patient admissions recorded in the NTDB. Of these, we excluded 563,326 whose payment mode was coded as no fault automobile, workers compensation, not billed, or other. Of the remaining 4,373,554 admissions, 953,281 (21.8%) were classified as uninsured and 3,420,273 (78.2%) as insured.

Figure 1 illustrates prevalence of each comorbid condition reported to the NTDB, stratified by insurance status. Most patients had at least one comorbidity (68.3%, n=2,985,089), with the most common conditions being hypertension (26.4%, n=1,153,404), “other” (19.1%, n=836,528), current smoker (13.1% n=572,255), and diabetes mellitus (10.2%, n=446,333). In general, insured patients were more likely to have any comorbidity (70.2 vs. 61.4%), as well as all comorbidities individually except current smoker (11.5 vs. 18.5%), alcoholism (6.0 vs. 10.0%), drug abuse or dependence (2.7 vs. 4.7%), and pre-hospital cardiac arrest (0.1 vs 0.3%).

**Figure 1.**
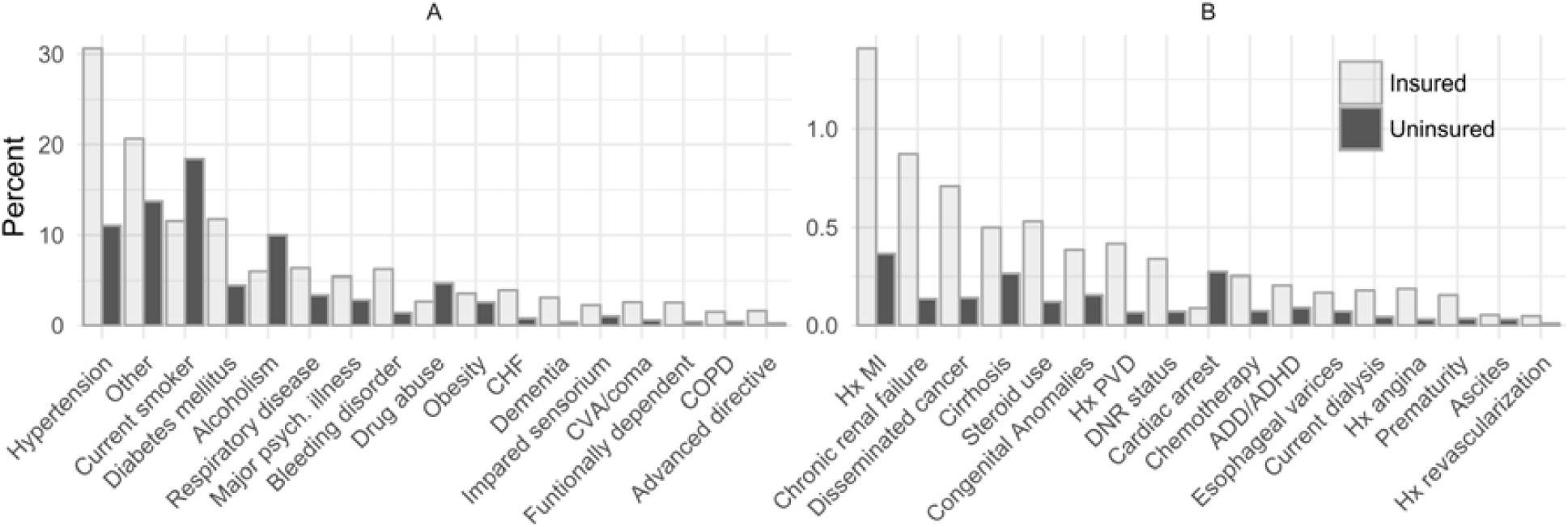
Prevalences of comorbid conditions in the National Trauma Data Bank, 2010-2015, separated by insurance status.

The scree plot depicting marginal proportion of variance explained by each additional axis in LPCA is displayed in Figure 2. Because an inflexion point occurred at 3 axes, after which each additional axis explains less than 7.5% of additional variance, we chose to include up to the first 3 principal axes in our models.

**Figure 2.**
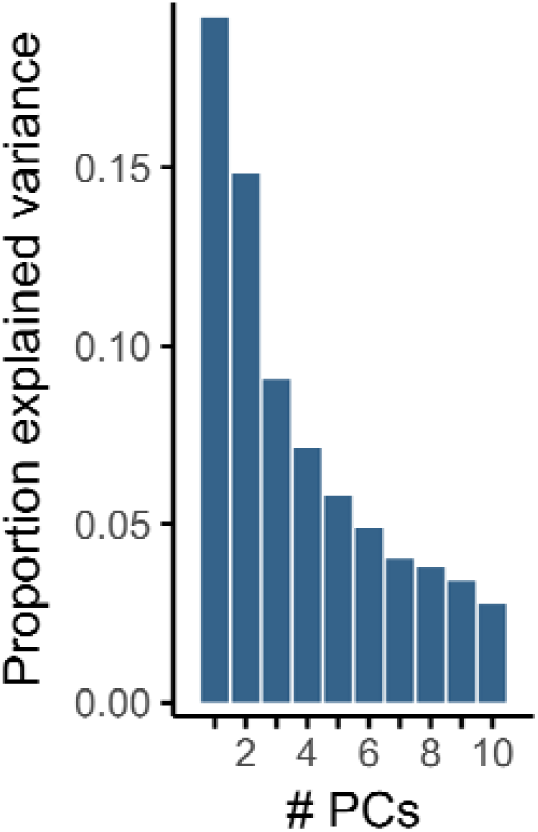
Marginal proportion of variance explained by each successive axis in logistic principal component analysis, calculated using a random sample of 50,000 without replacement. An inflection point exists at approximately 3 components, after which each additional component offers less than 7.5% increase in variance explained. Abbreviations: PCs, principal components.

Figure 3a shows to what extend each condition loads onto each of the first 3 principal axes, where the axes can be interpreted as propensities toward a particular type of patient. Patients with a high score for axis 1 would be likely to have hypertension, diabetes, as well as many of the other common conditions. In contrast, axis 2 describes the propensity to have hypertension and/or diabetes but no other common conditions except “other”, and axis 3 describes the propensity to have primarily hypertension and/or diabetes in the absence of other conditions. For example, a patient with hypertension, diabetes, bleeding disorder, CHF, and dementia has a score of 10.5 on axis 1 and 5.1 on axis 2, whereas a patient with only hypertension has a score −3.1 on axis 1 and 5.1 on axis 2. (For context, axis 1 ranges from −3.3 to 20.7, and axis 2 ranges from - 11.1 to 11.4.)

**Figure 3.**
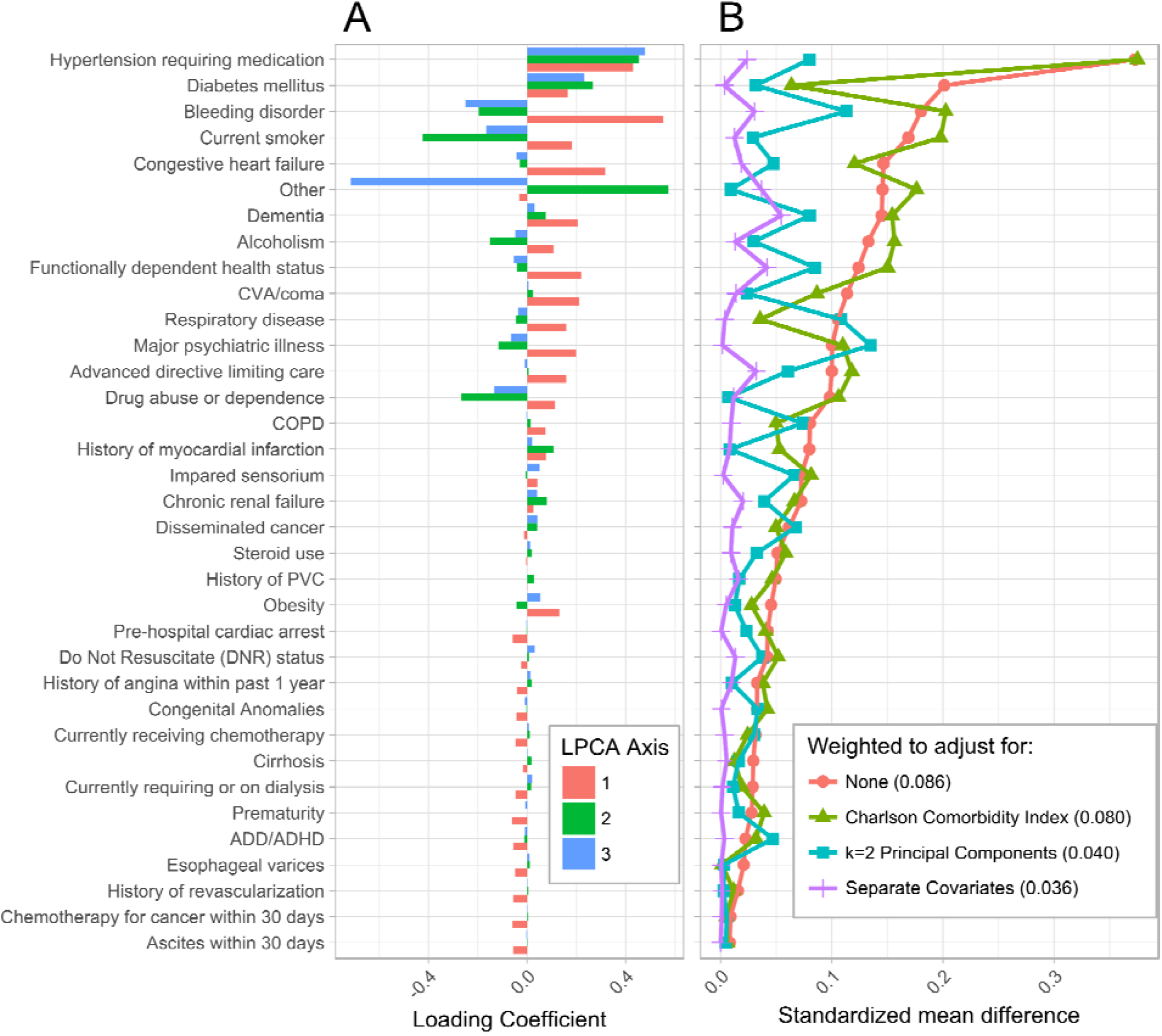
(A) Loading coefficients for the first three logistic principal components (LPCA). (B) Standardized mean differences (SMDs), both crude and weighted by IPTW. The sample weighted via the CCI performs similarly to the unweighted sample, while the sample weighted via the first 2 logistic principle components performs better, and the sample weighted via separate covariates performs the best, in terms of reduction in standardized mean differences. Numbers in parentheses in key are mean of SMDs.

Following calculation of IPTW weights based on CCI, LPCA, and each condition separately, we calculated crude and IPT-weighted standardized mean differences (SMDs) to evaluate covariate balance between insured and uninsured patients in crude vs. adjusted models, displayed in Figure 3b. Two metrics summarize the overall degree of covariate imbalance: the mean SMD and the percentage of SMDs which are greater than 0.1. While no consensus exists to our knowledge as to an acceptable cutoff for SMDs, 0.1 has been proposed as a rule of thumb for acceptable imbalance.^32^ In the unweighted sample, considerable imbalance exists (mean SMD=0.086, 37.8% >0.1). The sample weighted by CCI retains much of this imbalance (mean SMD=0.080, 32.4% >0.1), whereas the samples weighted by 2 (mean SMD 0.04, 8.1% >0.1) and 3 (mean SMD 0.036, 8.1% >0.1) LPCA axes retain little imbalance. Finally, as expected, the sample weighted by separate covariates has the least imbalance (mean SMD 0.012, 0% >0.1).

Following assessment of covariate balance using weighted SMDs, we fit logistic regression models, regressing probability of mortality on insurance status, both crude and adjusted separately for CCI, the first 2 and 3 LPCA axes, and comorbid conditions as separate covariates, using IPTW and standard regression adjustment (Table 2). In crude analysis, compared to insured patients, uninsured patients had 1.17 times the odds of mortality [95% CI 1.15 to 1.19]. After adjusting for each covariate separately, the odds were 1.36 times higher [95% CI 1.33 to 1.39] using IPTW and 1.4 times higher [95% CI 1.38 to 1.43] using regression adjustment. The estimates adjusting for the first two LPCA axes were attenuated slightly [OR_IPTW_ 1.31, 95% CI 1.26 to 1.32; OR_Regression_ 1.38, 95% CI 1.36 to 1.40] with respect to separate covariates. Adjusting for the CCI attenuated the estimates even further [OR_IPTW_ 1.25, 95% CI 1.22 to 1.28; OR_Regression_ 1.30, 95% CI 1.27 to 1.32]. Based on ORs derived from covariate-adjusted logistic regression, the CCI accounted for 56.1% of observed confounding, whereas the first 2 and 3 LPCA axes accounted for 91.3 and 94.6%, respectively.

**Table 2.**
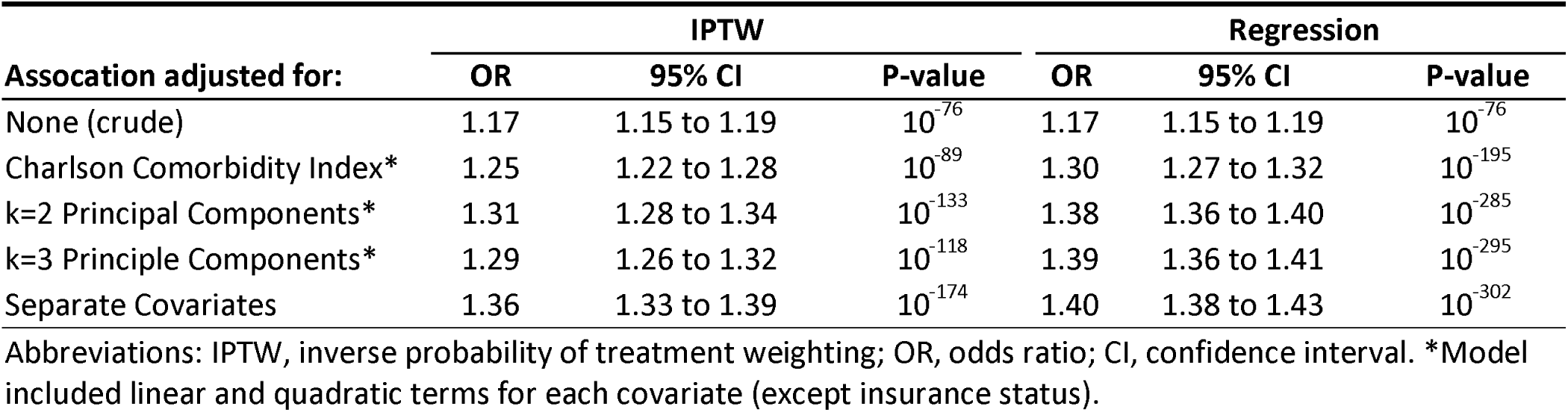
Crude and adjusted odds ratios for mortality in uninsured, compared to insured.

| Association adjusted for: | IPTW |  |  | Regression |  |  |
| --- | --- | --- | --- | --- | --- | --- |
|  | OR | 95% CI | P-value | OR | 95% CI | P-value |
| None (crude) | 1.17 | 1.15 to 1.19 | $10^{-76}$ | 1.17 | 1.15 to 1.19 | $10^{-76}$ |
| Charlson Comorbidity Index* | 1.25 | 1.22 to 1.28 | $10^{-89}$ | 1.30 | 1.27 to 1.32 | $10^{-195}$ |
| k=2 Principal Components* | 1.31 | 1.28 to 1.34 | $10^{-133}$ | 1.38 | 1.36 to 1.40 | $10^{-285}$ |
| k=3 Principle Components* | 1.29 | 1.26 to 1.32 | $10^{-118}$ | 1.39 | 1.36 to 1.41 | $10^{-295}$ |
| Separate Covariates | 1.36 | 1.33 to 1.39 | $10^{-174}$ | 1.40 | 1.38 to 1.43 | $10^{-302}$ |

Abbreviations: IPTW, inverse probability of treatment weighting; OR, odds ratio; CI, confidence interval. \*Model included linear and quadratic terms for each covariate (except insurance status).

## Discussion

This study investigated the properties of the Charlson Comorbidity Index (CCI), compared to logistic principal component analysis (LPCA) and separate comorbidity covariates, in terms of their ability to adequately control for measured confounding of the insurance-mortality relationship by comorbid conditions. We found that while adjusting for separate covariates induced excellent covariate balance, adjustment for the CCI did little to improve covariate balance beyond the crude analysis, whereas LPCA more closely approximated separate covariates. Adjusted estimates for the odds of mortality in uninsured vs. insured patients were also closer to the separate covariates estimates when using LPCA compared to CCI, both in IPTW and standard regression adjustment.

These results most likely reflect the limited definition of comorbidities reflected by the CCI – for instance, it does not include many common conditions such as hypertension, respiratory disease, and obesity. The score was developed using a backward stepwise Cox regression procedure in which conditions with mortality relative risk 1.2 or less were dropped,^3^ variables which could still be important confounders. In contrast, LPCA does not drop variables, but makes use of all available information. The differences in ORs derived from IPTW vs. regression adjustment likely reflect the non-collapsibility of the OR, as the former represents the marginal estimate and the latter the conditional estimate.^33^

These findings can help inform best practices for analysis of the NTDB, and potentially also other large inpatient databases. The CCI is widely used in practice to adjust for confounding by comorbidities, despite being developed solely to predict mortality. This practice may be a reflection of earlier works purporting to evaluate the ability of the CCI to adjust for confounding, which have stated that because the score had good predictive performance with respect to mortality and other outcomes (C-statistic = .77-.86), its use as a confounding control variable is warranted.^5, 6^ On the contrary, our study demonstrates that this practice is likely to result in considerable residual confounding, and therefore biased conclusions.

Based on our findings, the ideal method is to control for each covariate individually, but this practice becomes less feasible in smaller samples, as there may be a large number of potentially correlated comorbidity variables, along with violations of the positivity assumption. The use of a dimension reduction technique such as LPCA is well suited to this setting, as a very large number of potentially correlated variables can be compressed into an uncorrelated few, with most of the variation in the full dataset retained. Our findings demonstrate that LPCA, while less than ideal, will likely result in less residual confounding compared to a standardized index such as CCI.

Our findings should be interpreted in light of their limitations. As the NTDB is a voluntary registry with passive data collection, considerable measurement error may exist in comorbidity status; the methods presented herein are not intended to account for such error. We make the assumption that such measurement error is non-differential, and thus unlikely to affect the relative proportion of covariate imbalance accounted for by each method. Investigators will likely employ a variety of confounding control methods not limited to IPTW and regression adjustment, including propensity matching and stratification – while these should in principle produce similar results to ours, we cannot guarantee our results will generalize to these methods.

One limitation of LPCA is its requirement of considerable computational resources, compared to the simple arithmetic required to calculate the CCI, especially in very large datasets. However, the smaller datasets in which problems arise when controlling for ∼40 separate covariates are unlikely to present a computational problem for LPCA. Conversely, one possibility is for organizations compiling large data repositories to include in their public datasets a pre-calculated LPCA index of comorbidities, similar to the conditional mortality probability variables included in some datasets similar to the NTDB. Finally, this study is intended to be a methodological investigation and findings should not be interpreted scientifically – that is, there are likely a large number of additional confounders of the association between insurance and mortality, and our ORs may not be accurate effect estimates.

In conclusion, our findings suggest that the common practice of employing the CCI to adjust for confounding may result in residual confounding, and alternative strategies should be considered. While adjustment for each comorbidity individually appears to be the ideal method notwithstanding any measurement error in these variables, a dimensionality reduction method such as LPCA may be a viable alternative to adjusting for each comorbidity when samples are small or positivity assumptions are violated. Future research should aim to validate these findings in other datasets, in other research questions, and in other commonly used methods of covariate control such as propensity matching and stratification.

## Abbreviations

CCI: Charlson Comorbidity Index
PCA: principal components analysis
LPCA: logistic principal components analysis
IPTW: inverse probability of treatment weighting
SMD: standardized mean difference
NTDB: National Trauma Data Bank
ICD-9-CM: International Classification of Diseases, Ninth Revision, Clinical Modification
PVD: peripheral vascular disease
COPD: chronic obstructive pulmonary disorder
ADD/ADHD: attention deficit disorder/attention deficit hyperactivity disorder
ACS: American College of Surgeons
CHF: congestive heart failure

## Acknowledgments

The National Trauma Data Bank is full and copyrighted property of The American College of Surgeons, Committee on Trauma. The American College of Surgeons Committee on Trauma is not responsible for any claims arising from works based on the original data, text, tables, or figures herein.

Author contributions: AR and MB designed the study, AR carried out statistical analysis, AR composed the first draft of the manuscript, and AR and MB reviewed and edited the final manuscript.

## Notes

**Author Disclosure Statement** The authors have no conflicts of interest, financial or otherwise, to declare.

